# Boreal Forest Height Mapping using Sentinel-1 Time Series and improved LSTM model

**DOI:** 10.1101/2022.09.18.508417

**Authors:** Shaojia Ge, Hong Gu, Weimin Su, Yrjö Rauste, Jaan Praks, Oleg Antropov

## Abstract

Here, a novel semi-supervised Long Short-Term Memory (LSTM) model is developed and demonstrated for predicting forest tree height using time series of Sentinel-1 images. The model uses a Helix-Elapse (HE) projection approach to capture relationship between forest temporal patterns and Sentinel-1 time series, when the acquisition time intervals are irregular. A skip-link based LSTM block is introduced and a novel backbone network, Helix-LSTM, is proposed to retrieve temporal features at different receptive scales. Additionally, a novel semi-supervised strategy, Cross-Pseudo Regression, is employed to achieve better model performance. The developed model is compared versus basic LSTM model, attention-based bidirectional LSTM and several other established regression approaches used in forest variable mapping, demonstrating consistent improvement of forest height prediction accuracy. The study site is located in Central Finland and represents boreal forestland. At best, the achieved accuracy of forest height mapping was 28.3% rRMSE for pixel-level predictions, and 18.0% rRMSE on stand level. We expect that the developed model can also be used for modeling relationships between other forest variables and satellite image time series.

## I Introduction

**T**IMELY assessment and monitoring of forests forms basis for definition and implementation of preventive and corrective measures for sustainable forest management and forest restoration after disturbances [1]. Dynamics of forest structural variables provides information on forest status and forest changes and represents key information for forest management purposes [2], [3]. Traditional forest inventory variables, such as forest height, basal area, diameter at breast height and others can be used as inputs for biomass modelling. Furthermore, many users require information on traditional forest inventory variables as well, for example private forestry companies with smaller areas of interest, to support their forest management decisions.

Satellite based operational forest inventories often use satellite optical data augmented by reference plots for producing forest maps and estimates [4]. When use of optical satellite data is compromised due to near-permanent cloud coverage, possible solution is to use SAR sensors, relying on longer and denser time series. Radar based monitoring offers flexibility in forest monitoring applications, when results are requested in a fixed (e.g., yearly) schedule. Primary SAR data used in forest inventorying are L-band data because of smaller saturation of biomass-to-backscatter relationship compared to shorter wavelengths [5]–[7]. Use of advanced SAR temporal and textural features and imaging modes can improve accuracy of forest variable prediction [8], [9].

The European Copernicus program has opened new opportunities in forest mapping with the launch of Sentinel-1 satellites thanks to their high spatial and temporal resolution, ability to form long image time series, and data provision at no cost to users [10]. The Sentinel-1 mission consists of a constellation of two polar-orbiting satellites mounting a C-band synthetic aperture radar (SAR) imaging system. They offer a repeat cycle of six days with all-weather and day-and-night monitoring capabilities.

Multitemporal C-band SAR data were extensively used for evaluating and monitoring growing stock volume of both boreal and tropical forests, as well as in thematic mapping purposes [11]–[19]. Present consensus is that further research is required on methods exploiting dense time series of C-band SAR measurements, including multitemporal approaches, to achieve performance similar to L-band SAR data [7]. Important issues that need improvement is relatively poor prediction accuracy and lack of consistent ways to use EO time series data. One popular approach is to use multivariable models where each measurement/observation from time series is treated as independent feature. Such approaches are quite popular within machine learning and statistical non-parametric methodologies, and have been already demonstrated with Sentinel-1 time series in forest variable prediction and classification [9], [20], [21]. However, such approaches ignore temporal dependencies between the data. One of possible solutions to introduce a temporal context is to use LSTM models that can capture temporal relationships between images.

LSTMs were previously demonstrated in several remote sensing applications, particularly land cover mapping and crop monitoring [22]–[27]. To date, use of LSTMs in forest attribute prediction utilizing EO data was limited if at all reported with SAR image time series. Another general issue is lack of supervision data which could require semi-supervised approaches in model training. In this context, while consistency regularization has demonstrated success of semi-supervised learning algorithms in classification tasks [28], [29], there appears to be only few reports on its successful use in regression models for predicting continuous forest variables [30].

In this manuscript, we concentrate on use of LSTM based models for producing forest variable estimates from long time series of Sentinel-1 data. In particular we consider non-regular sampling of image datatakes and introduce a novel LSTM model that naturally takes varying time variable into account. We compare our approach with the classical LSTM model and several other pixel-based regression approaches that often used in satellite based forest inventory. We use forest tree height as a representative forest structural variable in our paper.

The paper is organized as follows. We describe our study site, acquired Sentinel-1 time series dataset and developed modeling approaches in Section II and Section III. Performance of developed model is analysed and compared to several benchmark approaches in Section IV, while potential challenges and opportunities are concluded in Section V.

## II Study Site, SAR and Reference Data

The study site featuring Hyytiälä forestry station is located in Central Finland, centre coordinates 37°2^*′*^*N*, 6°11^*′*^*E*. It is a square area covering 2500 km^2^. The location of the site is shown in Figure.1 along with an RGB color composite of three Sentinel-1 images. Typical boreal forest types are present in the study area, such as Norway spruce, Scots pine, and birch. Terrain is generally flat with the average elevation ranging from 95 m to 230 m above sea level. Frozen conditions often start in October in this area. The minimum temperature in winter can drop to − 25°C. First snow often falls in November and melts away completely by early May. The snow layer depth can reach from 20 cm to 70 cm depending on weather conditions.

Our dataset is represented by a long time series of Sentinel-1A IW-mode backscatter intensity images. Overall, there were 96 dual-polarization (VV+VH) images acquired from Oct. 9, 2014 to May 21, 2018. The time interval between adjacent observations ranges from 12 to 36 days. The original SAR data were acquired by Sentinel-1 satellites and initially pre-processed (focused and detected) and distributed by ESA as GRD (ground range detected) products. SAR image orthorectification and radiometric normalization was done using VTT in-house software [31]. Multilooking with factor of 2 × 2 (range × azimuth) was performed before the orthorectification. Radiometric normalization with respect to the projected area of the scattering element was performed to eliminate the topography-induced radiometric variation [32]. This way, a time series of co-registered “gamma-naught” backscatter images were formed, with a pixel size of 20 m by 20 m. Each image in the stack has size of 2500 px×2500 px corresponding to an area of 50 km×50 km.

Airborne laser scanning data (ALS) collected by National Land Survey of Finland in summer 2015 is used as reference data. The forest height was computed by averaging relative heights of ALS cloud points after the ground removal within each mapping unit. The height ranges from 0 m to 25.5 m and the average is 11.2 m. For additional comparisons with other conventional methods, stand-level estimates were also calculated from ALS data using a forest stand mask from Finnish Forest Centre.

We split the dataset and corresponding reference into three subsets: training, validation and testing as shown in Figure 1.c. Firstly, the whole area was equally divided into non-overlapping tiles, the size of each tile was 128 px × 128 px. Further, as depicted with red colour in Figure 2, 50% of the tiles were randomly selected, and pixels within them were extracted as the test subset. In a similar way, 10% of tiles were used to populate validation subset, and pixels from the remaining area composed the training dataset. The numbers of pixels are 1.5 mln, 0.375 mln and 1.778 mln for training, validation and testing subsets, respectively.

**Fig. 1.**
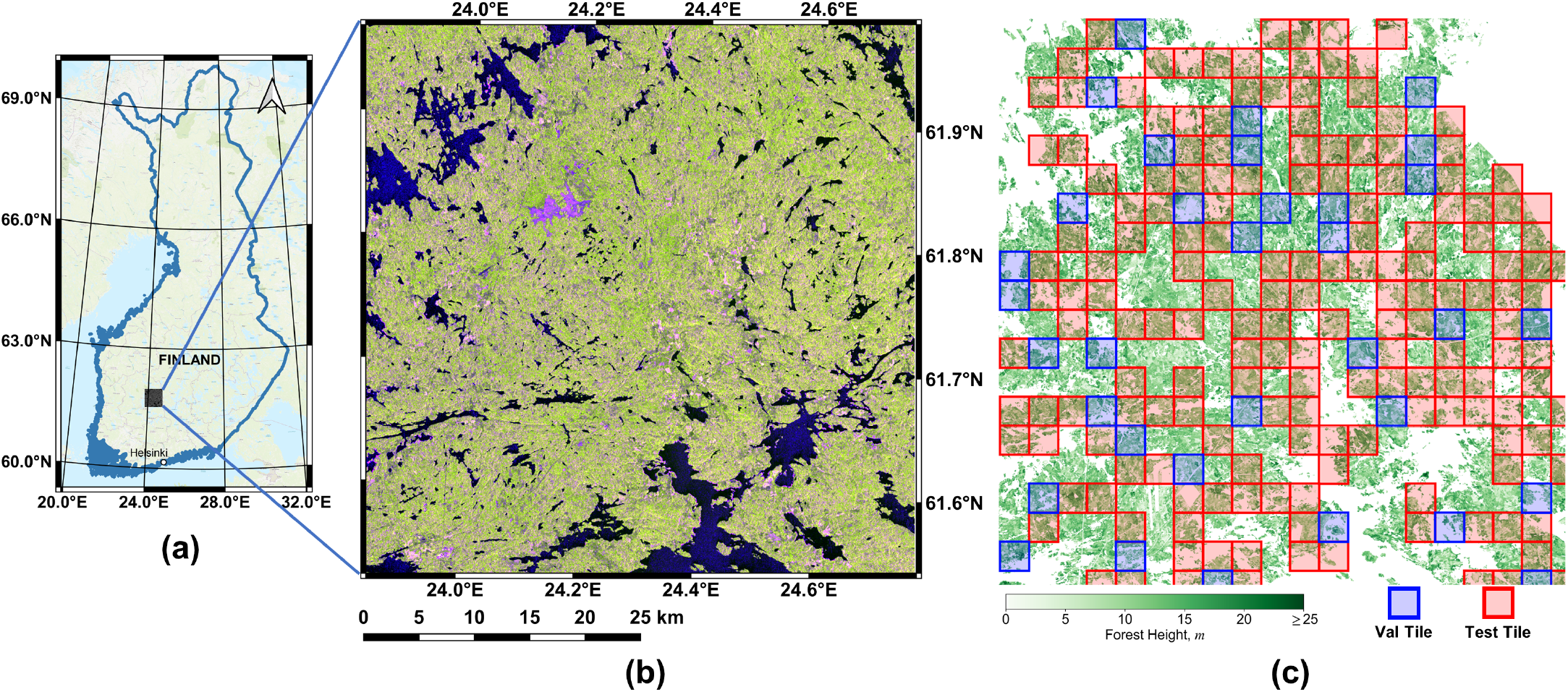
Study area: (a) location of study site in Finland, (b) RGB color-composite of 3 Sentinel-1 images, (c) reference ALS based forest height data with marked training and validation areas

**Fig. 2.**
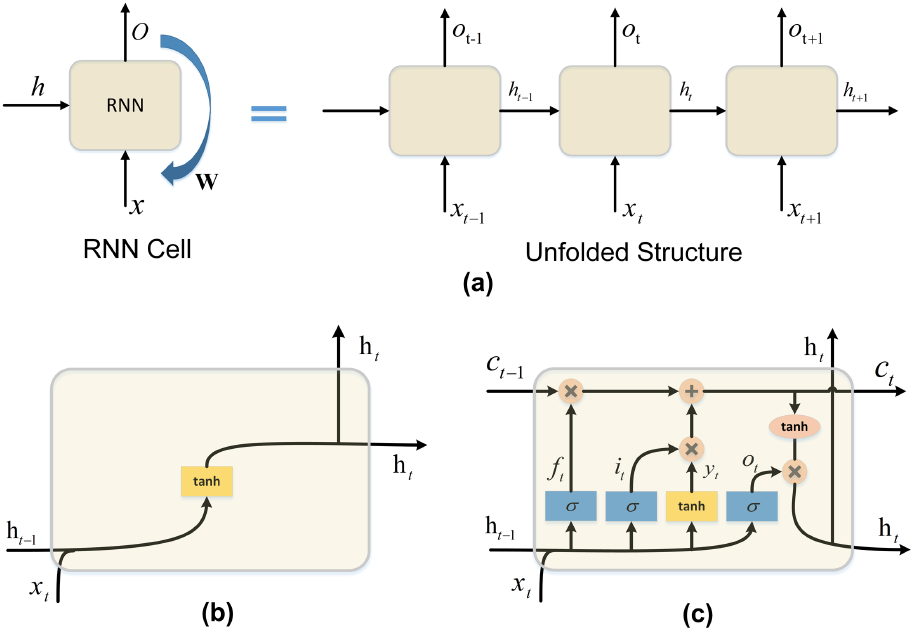
(a) The information flow diagram of RNN Cell (on the left) and its unfolded structure (on the right). The different two structures of (b) the classic RNN cell, and (c) LSTM cell.

## III Method Development

Here, we first briefly describe fundamentals of LSTMs and introduce a Helix-Elapse projection concept to deal with non-regular time interval between image datatakes. Further, we describe a semi-supervision regression and develop a model that combines several mentioned approaches. Baseline models that will be used in the benchmarking are also briefly described in this section.

### A. Long Short-Term Memory Networks

Contrary to convolutional neural network (CNN) [33], recurrent neural network (RNN) can clearly capture the temporal dependencies from a time sequence [34] thanks to its memory structure. However, classic RNN structures suffer from gradient vanishing or explosion problems and fail to capture long-term dependencies. To handle this issue, a revised cell structure was proposed in [35]. As shown in the diagram in Figure.2(c), the structure can be described as the “remember or forget” mechanism. Compared to the classical RNN cell, not only previous hidden state ***h***_***t*−1**_ but also previous memory ***c***_***t*−1**_ are fed into the current LSTM cell at timestamp *t*. They bring the history from previous timestamps into current workflow. Then by integrating three gates (input gate ***i***_***t***_, forget gate ***f***_***t***_ and output gate ***o***_***t***_), the flow of information are regulated and fused, and a decision on whether to keep it is performed using activation functions *tanh*. Finally, new current states, ***c***_***t***_ and ***h***_***t***_, are calculated and fed forward into the next timestamp.

The corresponding equations are mathematically described as

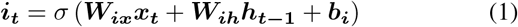

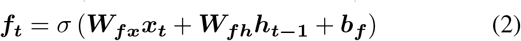

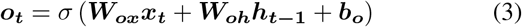

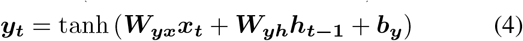

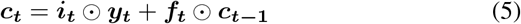

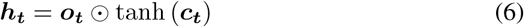

where *σ* and *tanh* represent Sigmoid and Hyperbolic activation function separately, ⊙ denotes the pointwise multiplication, ***W*** and ***b*** are weight matrix and bias vector.

### B. Helix-Elapse Projection

Timestamps of acquisitions contain seasonal information, and have explicit correlations with SAR image features, which may be considered as a prior knowledge for modeling. Especially, since time series with irregular timestamps often happen in real world practise, for instance, acquisition time interval differs from 12 to 36 in our case. How to alleviate the effect of irregular timestamps is an urgent problem. With timestamps as attributes, we can explicitly denote the acquisition interval differences, thus guide the training of the model.

We have already known that seasonal pattern apparently exists in forest remote sensing images. Other than just bringing raw timestamps as linear attributes, we map them into a two-dimensional space, where the transformed timestamps can form a helix curve. The mapping is done by a Helix-Elapse (HE) projection module, which can be mathematically described in Equation

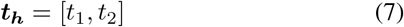

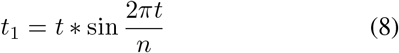

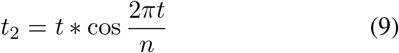

where *t* is the day index since Jan. 1st 2014, *n* is the total number of the days in a year, here we simply assign it as 365, ***t***_***h***_ is the projected timestamp vector.

As visualized in Figure 3, let the origin point be the start date (Jan. 1st 2014), the circulating angle of the helix curve indicates the date of a year, and the diameter indicates the growing year. The projected timestamps thus simulate the circulation of seasons in a heuristic way, as well as the year growing. We embed this HE projection module into our model, and then stack the output, helix time attributes ***T***_***h***_ = [***T***_**1**_, ***T***_**2**_], as two additional vectors together with the original input.

**Fig. 3.**
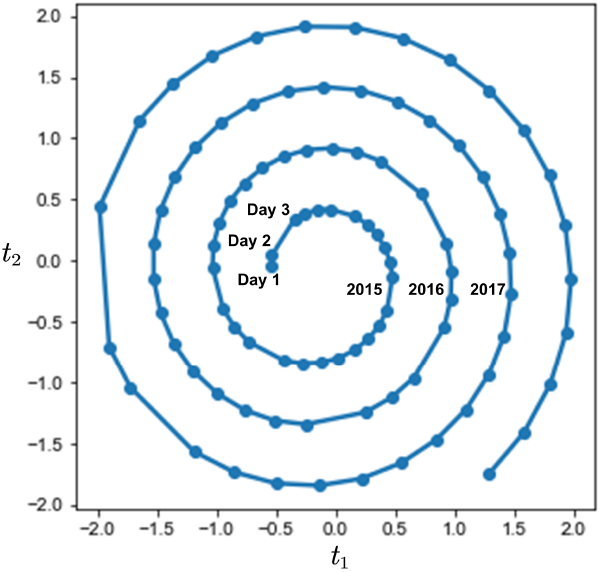
Helix time attributes for Sentinel-1 time series studied in the paper

### C. Skip-LSTM

In order to capture temporal dependencies within the SAR time series, we introduce LSTM as the backbone of our model. Despite carefully designed memory structure, LSTM in practice often fails to capture very long-term correlations [36]. In the case of satellite observations spanning several years, the time series would be particularly long. This type of long-term dependencies can hardly be captured by off-the-shelf recurrent units. Inspired by Dilated Convolution in CNN [37] and Recurrent-skip in LSTNet [36], we embed a Skip-LSTM module to alleviate the long-term correlation issues. The structure of Skip-LSTM is shown in Figure 4. It consists of two parts: convolutional layer and Skip-LSTM layer.

**Fig. 4.**
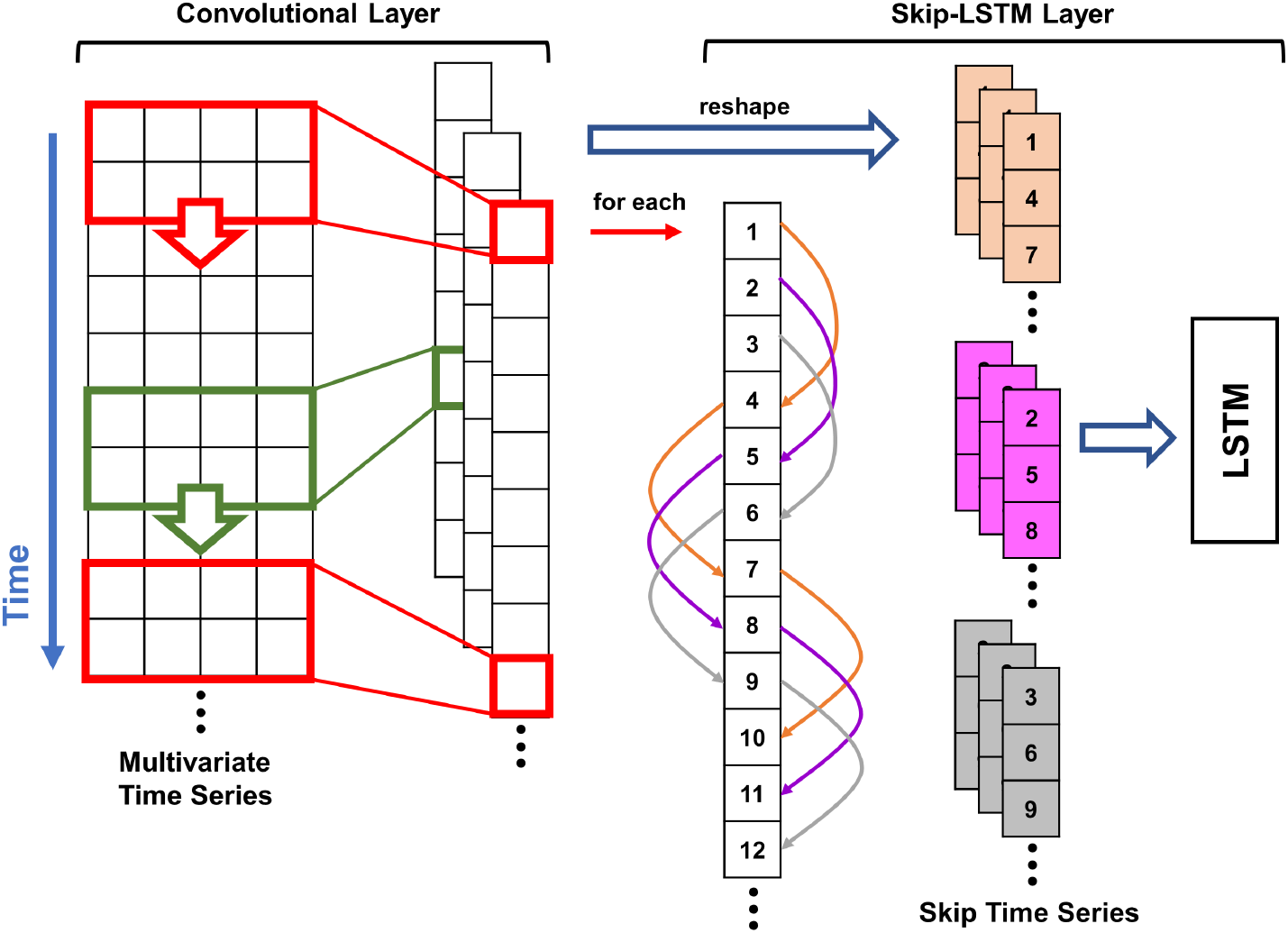
The structure of Skip-LSTM. The model consists of two layers: convolutional layer to fuse local temporal features, and Skip-LSTM layer to extract large-scale temporal dependencies.

Consider the input time series ***X*** with a size of *T* × *N*_*f*_, where the length of time series is *T* and the number of input channels is *N*_*f*_. A convolutional layer is first applied to capture short-term features in temporal domain as well as possible correlations between input channels. The convolutional layer is composed by multiple convolutional filters sweeping through the whole time series and extracting short-term features from the original input. The kernel size of each filter is *h* × *w*, where *h* decides the range of short-term features and *w* equals to *N*_*f*_. After the ReLU activation, the output has a size of *T* × *N*_*k*_, where *N*_*k*_ is the number of filters.

Further, a Skip-LSTM layer is applied on the product of convolutional layer. As shown in Figure 4, the skip-link structure of Skip-LSTM can jump over multiple timestamps, thus shortening the length of the time series. Following the skip-link, the raw time series are converted into a group of skip time series. Within each skip time series, non-adjacent timestamps get appended together. The total number of the converted series depends on the skip-factor *s*. Then the new grouped time series are fed into the backbone LSTM module as the input. By means of this “dilated receptive field”, long-term periodic patterns are captured in temporal domain at a larger scale.

### D. Cross-Pseudo Regression

A novel Cross-Pseudo regression (CPR) strategy [38] is further converted to wall-to-wall regression task to allow training the model in a semi-supervised way to compensate possible lack of training data. Both labeled and unlabeled data are included in the strategy. The model is constructed using two branches with the same structure but initialized differently, as shown in the flowchart of Figure 5.

**Fig. 5.**
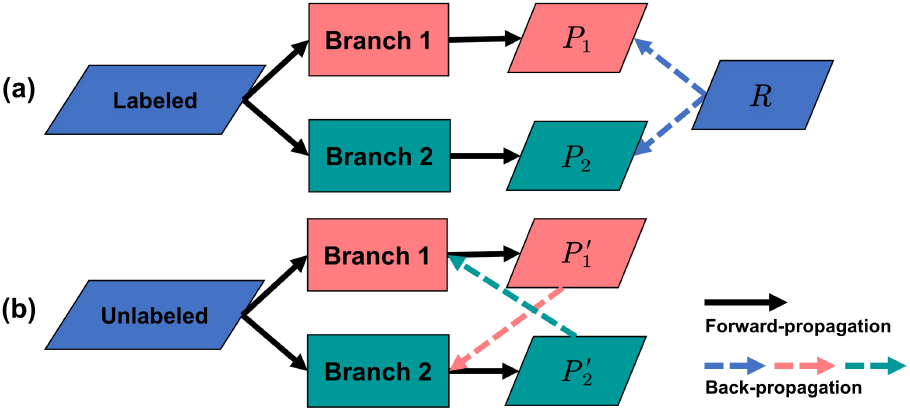
Cross-Pseudo regression strategy. (a) is the supervised step, (b) is the unsupervised step.

Firstly, we train the two branches separately in a normal supervised way, as shown in Figure 5.a, only labeled data is taken into the training. Specifically, we select mean squared error (MSE) to measure the distance between the predictions and reference, as mathematically described in Equation

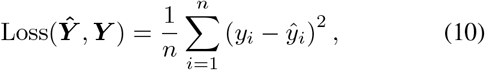

where ***Ŷ*** denotes the prediction and ***Y*** denotes the reference, *n* is the total number of the samples. So the supervised-loss can be denoted as

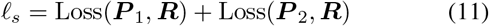

where ***P*** _1_ and ***P*** _2_ denote the predictions of Branch #1 and Branch #2 respectively, and ***R*** represents the reference.

Secondly, we ignore the label information and treat all the data as unlabeled, as illustrated in Figure 5.b. The results predicted by one branch can be naturally treated as pseudo-labels of the other, instead of the real reference. The back-propagation process can still be carried out according to the cross-pseudo-loss, which is defined as

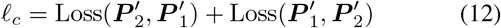

where 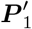 and 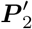 denote the pseudo labels of Branch #1 and Branch #2.

From the viewpoint of consistency regularization [39], [40], different initialization of the branches would bring perturbations to the model. When fed in the same input, both branches are encouraged to predict the same results, even though the perturbation is imposed. By minimizing the cross-pseudo loss, the discrepancy between predictions of both branches would also get minimized. By this way, a more compatible and representative feature space is learned by the model.

Finally, we combine supervised-loss and cross-pseudo-loss together. The combined loss of CPR can be defined as

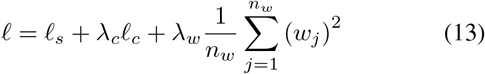

where *λ*_*c*_ controls the contribution of cross-pseudo loss, which is simply set as 0.5 in our study. *l*_2_ regularization item is added to the final loss to alleviate the overfitting, which is also known as the weight decay. *w*_*j*_ denotes the *j*-th weight in the model, *n*_*w*_ is the total number of the weights and *λ*_*w*_ decides the trade-off. On prediction stage, we simply select one of the trained branches, immigrate all its weights to task model, and give the final regression of the forest height.

### E. Overall Structure of CrsHelix-LSTM model

We name the proposed Cross Helix LSTM method as CrsHelix-LSTM for short. The overall schematic flowchart of the proposed architecture is shown in Figure. 6. The model consists of two branches based on CPR strategy. In each branch, helix time attributes are firstly mapped and attached to the input data. Then, the input is fed into two separate components to obtain comprehensive features in temporal domain. Vanilla LSTM is used to extract the overall invariant features of the time series. Convolutional layer is utilized to extract short-term features, followed by a Skip-LSTM layer to capture the seasonal pattern at a larger scope. At last, the features maps from both sides are stacked together. Regression header finally project the features to the forest height.

**Fig. 6.**
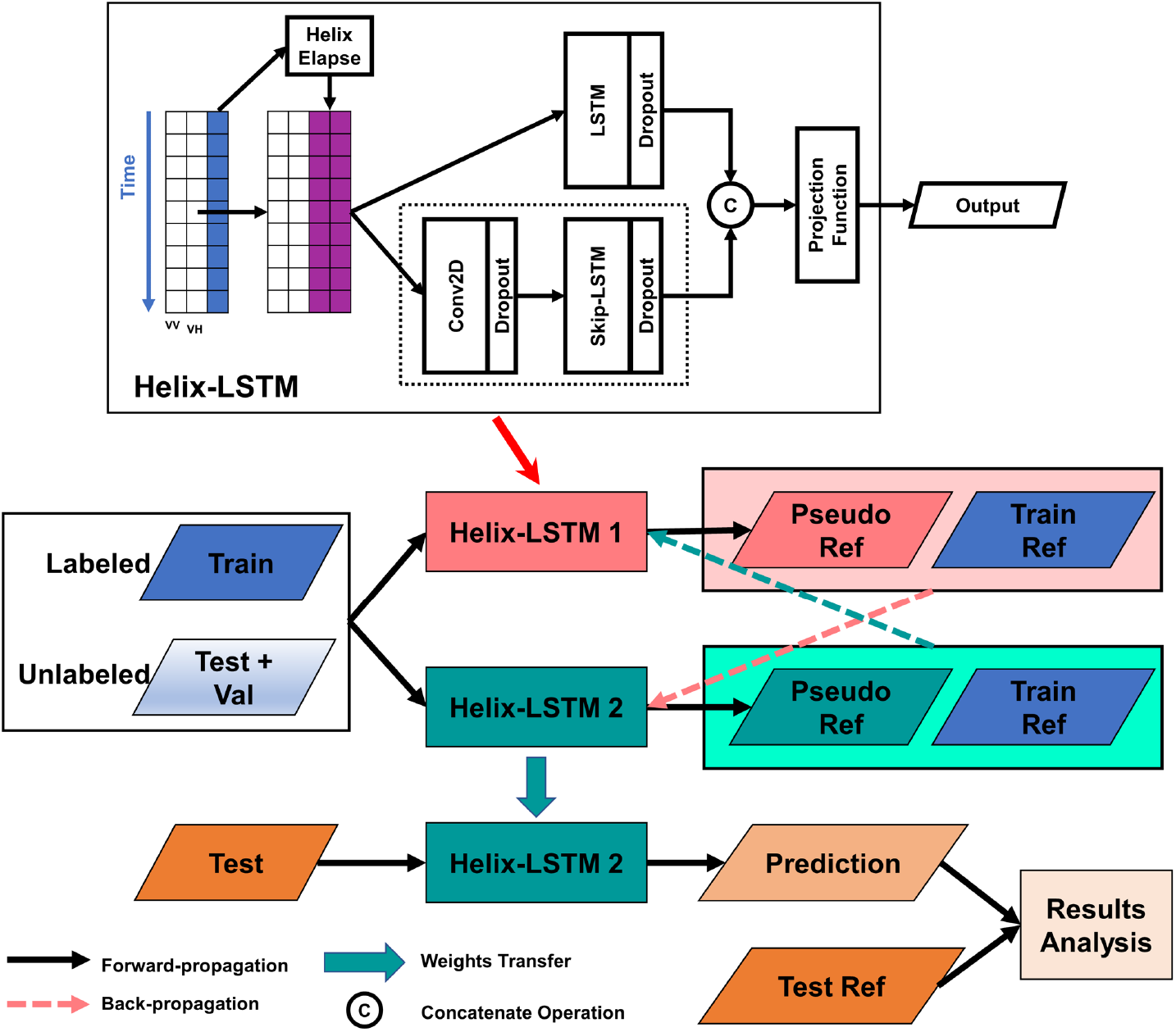
The overall flowchart of the proposed LSTM classification architecture.

The hidden size of LSTM and Skip-LSTM is set to 128, and for convolutional layer to 64. The kernel size of CNN module is 5, the skip-factor is set as 12.The total time step is 96, equivalent to the length of Sentinel-1 backscatter time series. To avoid potential overfitting, *Dropout* layers are implemented before regression header. The *Dropout* coefficient is set to 0.5.

At the training stage, we used the *Adam* optimisation algorithm to minimise the loss function. The batch size is 2048. *OneCycleLR* learning rate strategy took care of the training progress in case of overfitting [41].

### F. Baseline Models

To access added value of our proposed approach, several popular and more conventional regression methods were used as baselines for comparison in our study. It should be noted that, similar to our approach, all these methods operated on pixel-level considering only temporal features. Following modeling approaches are included for comparison:

- MLR, RF, LightGBM

Multiple Linear Regression (MLR) and Random Forest (RF) are mature regression methods, which have been widely used in forest remote sensing tasks. Light Gradient Boosting Machine (LightGBM) [42] is a modern Gradient Boosting Decision Tree (GBDT) model. Since proposed by Microsoft in 2017, its regression accuracy and efficiency has been proved in different application areas [43], [44].

- LSTM, Attn-BiLSTM

Since our proposed model is an improved version of LSTM, vanilla LSTM and its variant, Bidirectional LSTM with attention mechanism (Attn-BiLSTM) [45], [46], are also included as baseline models for comparison.

BiLSTM consists of two LSTMs with the same structure but opposite directions. Temporal dependencies are obtained from both directions. Furthermore, with the self-attention mechanism, the attention weights establish the correlations between timestamps, which reportedly can better tackle gradient vanishing problem and obtain long-term correlations [47]. Attn-BiLSTM combines these both merits, and has been introduced into some SAR remote sensing tasks.

To decrease the number of independent variables in MLR, a principal component analysis module was applied before the modeling. Such dimensionality reduction was not necessary for other methods, as they have built-in feature selection modules. All the models were fine-tuned with Optuna using 5-fold cross-validation. A 10% random samples of the training dataset was used in the fine-tuning.

### G. Model Prediction Accuracy Assessment

The prediction accuracy of various studies regression models was calculated using following accuracy metrics:

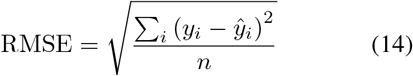

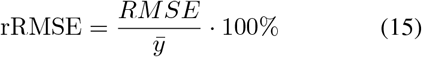

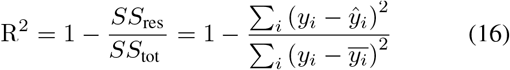

where *y*_*i*_ is the reference forest height of the pixel *i, Ŷ*_*i*_ is the predicted forest height, *n* is the total number of the samples. Stand-level estimates of forest height were calculated using spatial averaging at extent of each stand (available from forest stand mask), with reported accuracies calculated using exactly the same equations (14)-(18) for aggregated stand-level units.

## IV Experimental Results and Discussion

### A. Experimental Settings

The experiments were performed using Windows Server with Intel Xeon E5-2697 v4 CPU and NVIDIA GTX3060 GPU accelerated by CUDA 11.3 toolkit. LSTM, Attn-BiLSTM and the proposed model were built with a neural network library, Pytorch 1.11.0. MLR and RF were implemented with scikit-learn machine learning toolbox. LightGBM was implemented with LightGBM Python-package provided by Microsoft.

### B. General Performance Evaluation

We have examined performance of developed model versus several other LSTM approaches, as well as a representative set of benchmarking models often used in forest mapping, on both pixel-level and stand-level. Obtained ablation results for LSTM based approaches are gathered in Table I, and accuracy metrics for benchmarking models are shown in Table II. Examples of produced forest maps with various approaches are shown in Figure 7. Prediction performance are gathered and illustrated with scatterplots between predicted and reference forest height in Figure 8.

**TABLE I.**
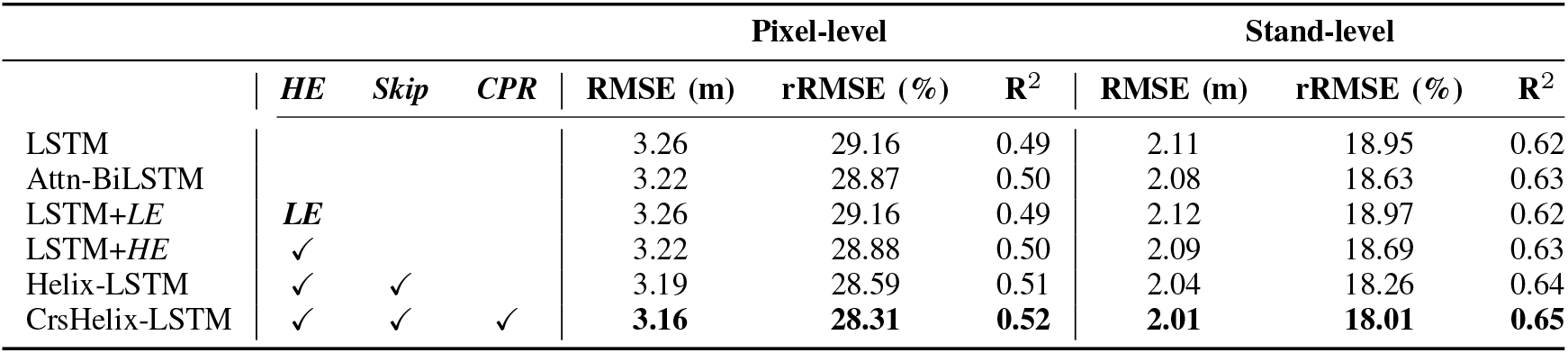
Ablation Study. *LE* stands for Linear-Elapse projection, *HE* for Helix-Elapse projection, *Skip* for Skip-LSTM block, and *CPR* for CPR strategy.

**TABLE II.**
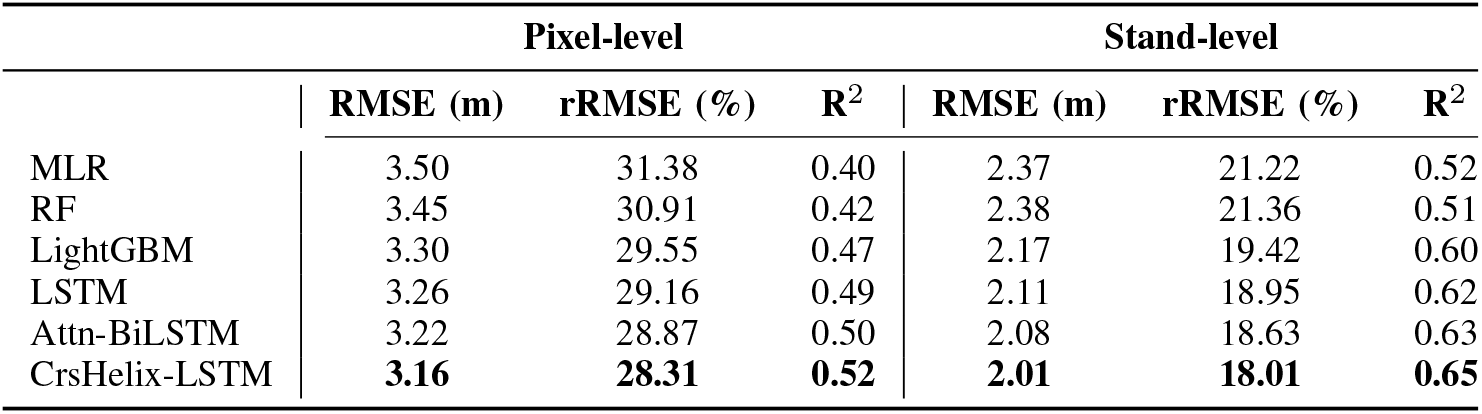
Experimental results compared to benchmarks

|  | Pixel-level |  |  | Stand-level |  |  |
| --- | --- | --- | --- | --- | --- | --- |
|  | RMSE (m) | rRMSE (%) | R<sup>2</sup> | RMSE (m) | rRMSE (%) | R<sup>2</sup> |
| MLR | 3.50 | 31.38 | 0.40 | 2.37 | 21.22 | 0.52 |
| RF | 3.45 | 30.91 | 0.42 | 2.38 | 21.36 | 0.51 |
| LightGBM | 3.30 | 29.55 | 0.47 | 2.17 | 19.42 | 0.60 |
| LSTM | 3.26 | 29.16 | 0.49 | 2.11 | 18.95 | 0.62 |
| Attn-BiLSTM | 3.22 | 28.87 | 0.50 | 2.08 | 18.63 | 0.63 |
| CrsHelix-LSTM | **3.16** | **28.31** | **0.52** | **2.01** | **18.01** | **0.65** |

**Fig. 7.**
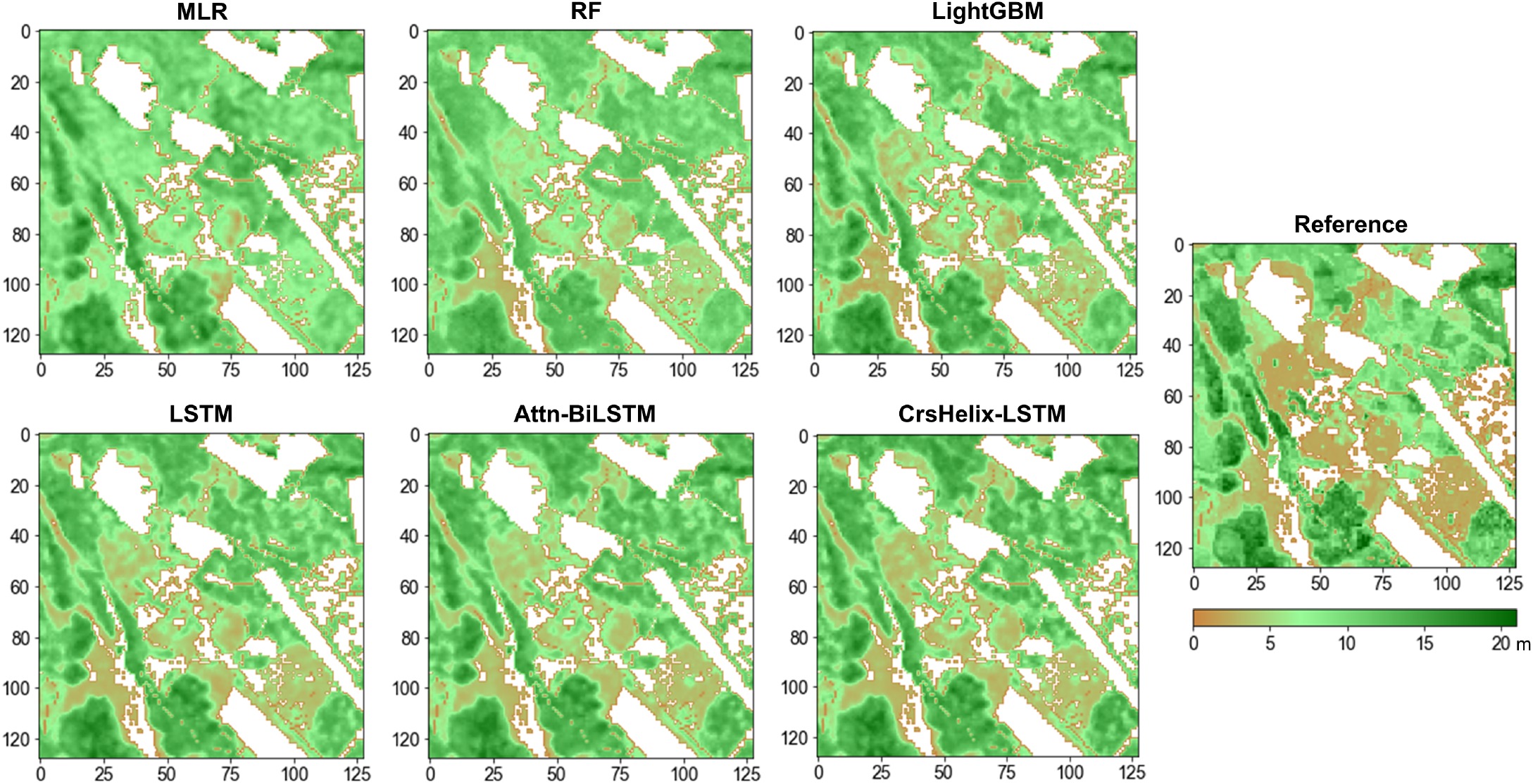
Examples of produced forest height maps for various examined regression approaches.

**Fig. 8.**
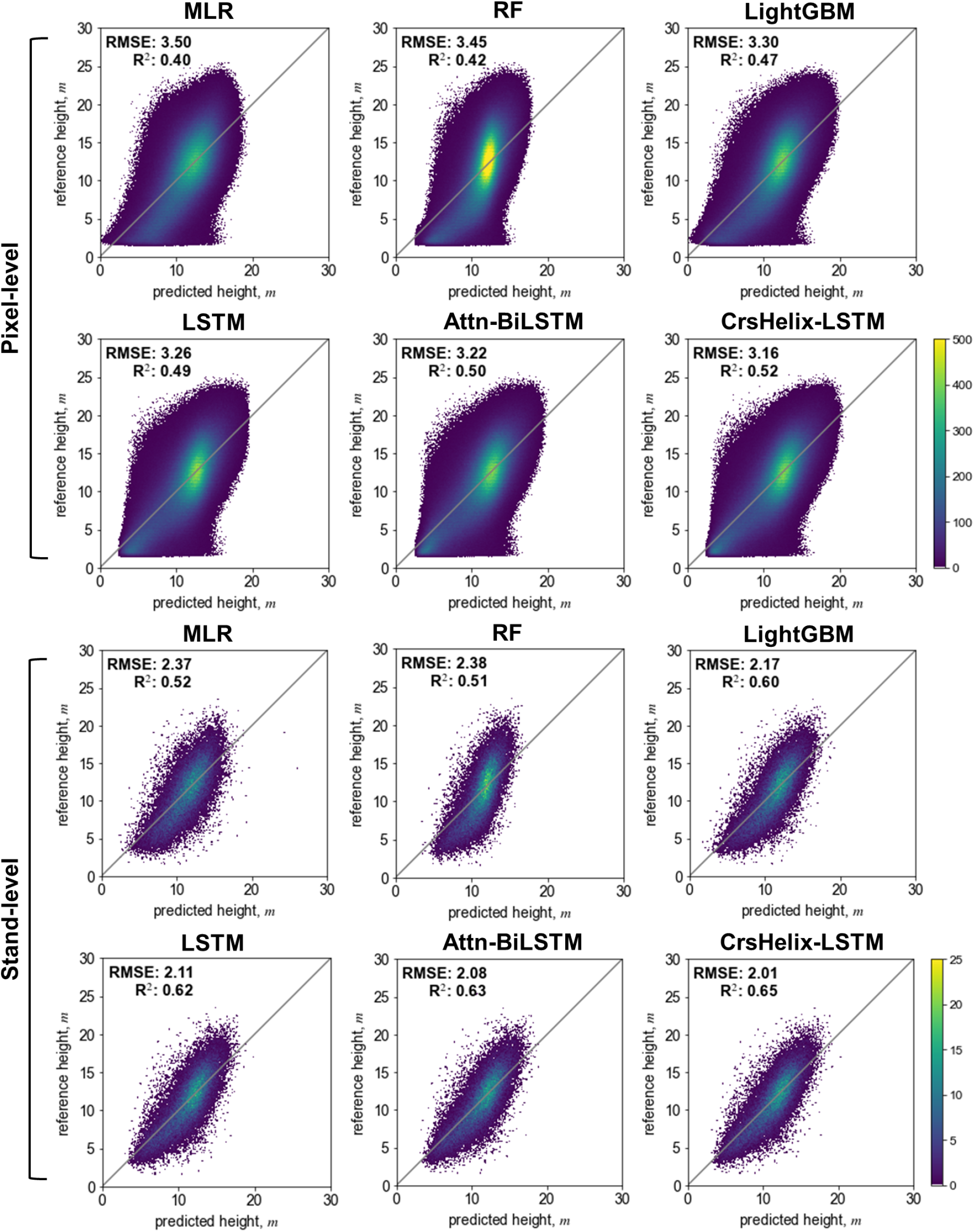
Pixel- (upper rows) and stand-level (bottom rows) scatterplots for various studied regression models: MLR, RF, LGBM, LSTM, Attn-BiLSTM with attention and CrsHelix-LSTM approach.

#### 1) Ablation Study

Firstly, we verified the effectiveness of different blocks in the ablation study, the results are shown in Table I. In general, forest height prediction at stand-level demonstrates larger accuracy compared to pixel-level, taking advantage from averaging within the homogeneous stands. When we handled the irregular time intervals by using Linear-Elapse (LE) projection, the LE attribute was stacked onto the input as a new feature. The regression results is not improved compared to Vanilla LSTM. It indicates the elapsing of time is not properly modeled in this case. On the contrary, by simply substituting the LE attributes with our Helix-Elapse (HE), RMSE is somewhat improved from 3.26 m to 3.22 m at pixel-level. The results are close to attention-based BiLSTM, which is considered as a much stronger baseline when it comes to long time series tasks. It indicates HE projection can better model the annual dynamics and help establish relationships between forest variables and seasonal patterns thereby approximating the role of attention mechanism at a lower computational cost.

When the Skip-LSTM block was embedded into the model, the regression performance of Helix-LSTM further improved at both pixel- and stand-level. The rRMSE is 28.59%, getting 0.28% improved from Attn-BiLSTM for pixel-level and 0.37% for stand-level. Considering Attn-BiLSTM is only better than Vanilla-LSTM for 0.29% and 0.32%, the improvement is quantitatively considerable. Finally, by wrapping up two Helix-LSTM branches with CPR strategy, CrsHelix-LSTM got the best regression performance in the ablation study. The best rRMSE is 28.31% for pixel-level and 18.01% for stand-level, and R^2^ of the final model is as high as 0.65. To note that the rRMSE gets 0.28% decreased further compared to Helix-LSTM, even though the backbone models were the same. It indicates an positive impact is imposed by the semi-supervised learning strategy, the model is learning the forest representations from not only the labeled but also unlabeled data.

#### 2) Method Performance Comparison with Baseline approaches

In Table II, we also compare our methods to some existing machine learning models. MLR, which is most widely used in forest mapping, presents the most fundamental performance that is 3.50 m of RMSE for pixel-level and 2.37 m for stand-level. Other approaches generally perform better than MLR, among them the use of LSTM somewhat improves the prediction performance at both pixel- and stand-levels. While the developed CrsHelix-LSTM approach has provided consistently better results than other benchmarked models. Its rRMSE for pixel-level reaches to 28.31%, 1.24% decreased compared to LightGBM. Note that rRMSE of LightGBM is only 1.83% decreased compared to MLR, from this aspect the improvement of our model is considerable.

Similar conclusions can also be drawn from analyses of scatter plots shown in Figure 8, particularly on stand-level. Compared to other benchmarked methods, the samples predicted by our CrsHelix-LSTM are more inclined to follow the diagonal line. Taller referenced samples around 20 m are less biased, which indicates a better prediction performance for taller forest stands. By visualizing the prediction results, as shown in Figure 7, we can better observe the prediction discrepancy over forests at different height levels. Compared to other methods, CrsHelix-LSTM is more sensitive to under-growth forests whose height is within 5 m. The corresponding areas are showing as light brown according to the colormap in Figure 7.

### C. Comparison with Other Studies and Outlook

Observed experimental results are encouraging further investigations and are generally in line with other reported studies in boreal forest biome [3], [7], [48], [49]. Obtained accuracies are notably higher than in several other studies using Sentinel-1 data or Sentinel-2 datasets or their combinations, and compare well versus earlier multisensor EO data studies [3], [20], [48]–[52]. Several datasets with high potential for forest variable retrieval, such as e.g. TanDEM-X, have relatively sparse coverage (both in geographic and temporal domain) and limited availability and thus not fully suitable suitable for large-area mapping and persistent monitoring purposes.

In boreal region, studies on forest variable prediction using Sentinel-1 or Landsat data report prediction accuracies within the range of 35-60% rRMSE [3], [48], while proposed model utilizing Sentinel-1 time series data reached rRMSE as small as 18 %. Predictions obtained using traditional ML models were within the same accuracy range as in recently published studies using Sentinel-2 and Landsat [53], while predictions using different versions of LSTM models appeared more accurate. Use of attention mechanism and Helix attribute was instrumental in providing larger accuracies of proposed Attn-LSTM and CrsHelix-LSTM models compared to basic LSTM model.

There is relatively limited literature using SAR data for forest height predictions, with predictions often reported on stand-level or coarser resolution spatial units [7]. However, our stand-level tree height predictions were at the same accuracy level or even better then reported retrievals with TanDEM-X interferometric SAR data deemed much more suitable for vertical forest structure retrieval [54]–[56]. To the best of our knowledge, obtained accuracy levels of 18% RMSE for boreal forest using CrsHelix-LSTM and combined SAR and optical data are superior to earlier results reported in literature [7].

## V Conclusions

Our study demonstrated the potential for applying LSTM approaches with Helix attribute for predicting forest structural variables such as forest height. A novel LSTM model incorporating Helix-Elapse projection, Skip-LSTM and Cross-Pseudo regression has been developed and tested in the study. The developed model was demonstrated using long time series of Sentinel-1 data but can be applicable to other EO data time series. The CrsHelix-LSTM model provided larger accuracies in boreal forest height mapping in Central Finland compared to other evaluated LSTM approaches and a set of representative machine learning approaches often used in forest mapping.

Future work will concentrate on introducing other datasets particularly suitable for retrieving vertical structure of forests, such as Sentinel-1 interferometric SAR and TanDEM-X datasets, as well as studying other forest variables, such as growing stock volume and above-ground biomass.

## Acknowledgments

This study was supported by the National Natural Science Foundation of China (Grant No. 61801221, 62001229, 62101264), and by China Postdoctoral Science Foundation (Grant No. 2020M681604). O.A. was supported by Multico project funded by Business Finland and Forest Carbon Monitoring project funded by European Space Agency.

## References

[1] M. Herold, S. Carter, V. Avitabile, A. B. Espejo, I. Jonckheere, R. Lucas, R. E. McRoberts, E. Næsset, J. Nightingale, R. Petersen, J. Reiche, E. Romijn, A. Rosenqvist, D. M. A. Rozendaal, F. M. Seifert, M. J. Sanz, and V. De Sy, “The role and need for space-based forest biomass-related measurements in environmental management and policy,” Surveys in Geophysics, vol. 40, no. 4, p. 757–778, 2019, cited by: 40; All Open Access, Green Open Access, Hybrid Gold Open Access. [Online]. Available: https://www.scopus.com/inward/record.uri?eid=2-s2.0-85061374917&doi=10.1007%2fs10712-019-09510-6&partnerID=40&md5=04cb405621e5c6468ee95d498f09fd1c

[2] R. E. McRoberts and E. O. Tomppo, “Remote sensing support for national forest inventories,” Remote Sensing of Environment, vol. 110, no. 4, pp. 412–419, 2007, forestSAT Special Issue.

[3] J. Miettinen, S. Carlier, L. Häme, A. Mäkelä, F. Minunno, J. Penttilä, J. Pisl, J. Rasinmäki, Y. Rauste, L. Seitsonen, X. Tian, and T. Häme, “Demonstration of large area forest volume and primary production estimation approach based on sentinel-2 imagery and process based ecosystem modelling,” International Journal of Remote Sensing, vol. 42, no. 24, p. 9492–9514, 2021.

[4] E. Tomppo, H. Olsson, G. Ståhl, M. Nilsson, O. Hagner, and M. Katila, “Combining national forest inventory field plots and remote sensing data for forest databases,” Remote Sensing of Environment, vol. 112, no. 5, pp. 1982–1999, 2008, earth Observations for Terrestrial Biodiversity and Ecosystems Special Issue. [Online]. Available: http://www.sciencedirect.com/science/article/pii/S0034425708000242

[5] T. Le Toan, A. Beaudoin, J. Riom, and D. Guyon, “Relating forest biomass to sar data,” IEEE Transactions on Geoscience and Remote Sensing, vol. 30, no. 2, pp. 403–411, 1992.

[6] M. L. Imhoff, “Radar backscatter and biomass saturation: ramifications for global biomass inventory,” IEEE Transactions on Geoscience and Remote Sensing, vol. 33, no. 2, pp. 511–518, 1995.

[7] GFOI, Integrating remote-sensing and ground-based observations for estimation of emissions and removals of greenhouse gases in forests: Methods and Guidance from the Global Forest Observations Initiative. Pub: Group on Earth Observations, Geneva, Switzerland, 2014.

[8] C. Schmullius, C. Thiel, C. Pathe, and M. Santoro, “Radar time series for land cover and forest mapping,” in Remote Sensing Time Series. Springer, 2015, pp. 323–356.

[9] E. Tomppo, O. Antropov, and J. Praks, “Boreal forest snow damage mapping using multi-temporal sentinel-1 data,” Remote Sensing, vol. 11, no. 4, 2019. [Online]. Available: https://www.mdpi.com/2072-4292/11/4/384

[10] R. Torres, P. Snoeij, D. Geudtner, D. Bibby, M. Davidson, E. Attema, P. Potin, B. Rommen, N. Floury, M. Brown et al., “Gmes sentinel-1 mission,” Remote Sensing of Environment, vol. 120, pp. 9–24, 2012.

[11] C. Thiel, O. Cartus, R. Eckardt, N. Richter, C. Thiel, and C. Schmullius, “Analysis of multi-temporal land observation at c-band,” in 2009 IEEE International Geoscience and Remote Sensing Symposium, vol. 3, 2009, pp. III–318–III–321.

[12] O. Antropov, Y. Rauste, A. Väänänen, T. Mutanen, and T. Häme, “Mapping forest disturbance using long time series of sentinel-1 data: Case studies over boreal and tropical forests,” in 2016 IEEE International Geoscience and Remote Sensing Symposium (IGARSS), 2016, pp. 3906–3909.

[13] G. V. Laurin, J. Balling, P. Corona, W. Mattioli, D. Papale, N. Puletti, M. Rizzo, J. Truckenbrodt, and M. Urban, “Above-ground biomass prediction by Sentinel-1 multitemporal data in central Italy with integration of ALOS2 and Sentinel-2 data,” Journal of Applied Remote Sensing, vol. 12, 2018.

[14] M. A. Stelmaszczuk-Górska, M. Urbazaev, C. Schmullius, and C. Thiel, “Estimation of above-ground biomass over boreal forests in Siberia using updated in situ, ALOS-2 PALSAR-2, and RADARSAT-2 data,” Remote Sensing, vol. 10, no. 10, 2018.

[15] O. Antropov, Y. Rauste, J. Praks, F. M. Seifert, and T. Häme, “Mapping forest disturbance due to selective logging in the congo basin with radarsat-2 time series,” Remote Sensing, vol. 13, no. 4, 2021. [Online]. Available: https://www.mdpi.com/2072-4292/13/4/740

[16] E. Tomppo, G. Ronoud, O. Antropov, H. Hytönen, and J. Praks, “Detection of forest windstorm damages with multitemporal sar data—a case study: Finland,” Remote Sensing, vol. 13, no. 3, 2021. [Online]. Available: https://www.mdpi.com/2072-4292/13/3/383

[17] M. Rüetschi, D. Small, and L. T. Waser, “Rapid detection of windthrows using sentinel-1 c-band sar data,” Remote Sensing, vol. 11, no. 2, 2019.

[18] D. Hoekman, B. Kooij, M. Quiñones, S. Vellekoop, I. Carolita, S. Budhiman, R. Arief, and O. Roswintiarti, “Wide-area near-real-time monitoring of tropical forest degradation and deforestation using sentinel-1,” Remote Sensing, vol. 12, no. 19, 2020. [Online]. Available: https://www.mdpi.com/2072-4292/12/19/3263

[19] M. G. Hethcoat, J. M. Carreiras, D. P. Edwards, R. G. Bryant, and S. Quegan, “Detecting tropical selective logging with c-band sar data may require a time series approach,” Remote Sensing of Environment, vol. 259, p. 112411, 2021. [Online]. Available: https://www.sciencedirect.com/science/article/pii/S0034425721001292

[20] S. Ge, E. Tomppo, Y. Rauste, R. E. McRoberts, J. Praks, H. Gu, W. Su, and O. Antropov, “Using hypertemporal Sentinel-1 data to predict forest growing stock volume,” bioRxiv, 2021.

[21] A. Dostálová, W. Wagner, M. Milenković, and M. Hollaus, “Annual seasonality in sentinel-1 signal for forest mapping and forest type classification,” International Journal of Remote Sensing, vol. 39, no. 21, pp. 7738–7760, 2018.

[22] S. Ge, O. Antropov, W. Su, H. Gu, and J. Praks, “Deep recurrent neural networks for land-cover classification using sentinel-1 insar time series,” in IGARSS 2019 - 2019 IEEE International Geoscience and Remote Sensing Symposium, 2019, pp. 473–476.

[23] D. Ienco, R. Gaetano, C. Dupaquier, and P. Maurel, “Land cover classification via multitemporal spatial data by deep recurrent neural networks,” IEEE Geoscience and Remote Sensing Letters, vol. 14, no. 10, pp. 1685–1689, 2017.

[24] Y. Yuan, L. Lin, L.-Z. Huo, Y.-L. Kong, Z.-G. Zhou, B. Wu, and Y. Jia, “Using an attention-based lstm encoder–decoder network for near real-time disturbance detection,” IEEE Journal of Selected Topics in Applied Earth Observations and Remote Sensing, vol. 13, pp. 1819–1832, 2020.

[25] L. Ma, Y. Liu, X. Zhang, Y. Ye, G. Yin, and B. A. Johnson, “Deep learning in remote sensing applications: A meta-analysis and review,” ISPRS Journal of Photogrammetry and Remote Sensing, vol. 152, pp. 166–177, 2019. [Online]. Available: https://www.sciencedirect.com/science/article/pii/S0924271619301108

[26] Y. Xie and J. Huang, “Integration of a crop growth model and deep learning methods to improve satellite-based yield estimation of winter wheat in henan province, china,” Remote Sensing, vol. 13, no. 21, 2021. [Online]. Available: https://www.mdpi.com/2072-4292/13/21/4372

[27] W. L. Hakim, A. S. Nur, F. Rezaie, M. Panahi, C.-W. Lee, and S. Lee, “Convolutional neural network and long short-term memory algorithms for groundwater potential mapping in anseong, south korea,” Journal of Hydrology: Regional Studies, vol. 39, p. 100990, 2022.

[28] Y. Wang, C. M. Albrecht, N. A. A. Braham, L. Mou, and X. X. Zhu, “Self-supervised learning in remote sensing: A review,” arXiv preprint arXiv:2206.13188, 2022.

[29] C. Wang, H. Gu, and W. Su, “Sar image classification using contrastive learning and pseudo-labels with limited data,” IEEE Geoscience and Remote Sensing Letters, vol. 19, pp. 1–5, 2021.

[30] S. Ge, H. Gu, W. Su, J. Praks, and O. Antropov, “Improved semisupervised unet deep learning model for forest height mapping with satellite sar and optical data,” IEEE Journal of Selected Topics in Applied Earth Observations and Remote Sensing, vol. 15, pp. 5776–5787, 2022.

[31] Y. Rauste, A. Lonnqvist, M. Molinier, J.-B. Henry, and T. Hame, “Orthorectification and terrain correction of polarimetric sar data applied in the alos/palsar context,” in 2007 IEEE International Geoscience and Remote Sensing Symposium. IEEE, 2007, pp. 1618–1621.

[32] D. Small, “Flattening gamma: Radiometric terrain correction for sar imagery,” IEEE Transactions on Geoscience and Remote Sensing, vol. 49, no. 8, pp. 3081–3093, 2011.

[33] A. Krizhevsky, I. Sutskever, and G. E. Hinton, “Imagenet classification with deep convolutional neural networks,” Advances in neural information processing systems, vol. 25, 2012.

[34] A. Graves, A.-R. Mohamed, and G. Hinton, “Speech recognition with deep recurrent neural networks,” in IEEE ICASSP Proc. IEEE, 2013, pp. 6645–6649.

[35] S. Hochreiter and J. Schmidhuber, “Long short-term memory,” Neural computation, vol. 9, no. 8, pp. 1735–1780, 1997.

[36] G. Lai, W.-C. Chang, Y. Yang, and H. Liu, “Modeling long-and short-term temporal patterns with deep neural networks,” in The 41st International ACM SIGIR Conference on Research & Development in Information Retrieval, 2018, pp. 95–104.

[37] F. Yu and V. Koltun, “Multi-scale context aggregation by dilated convolutions,” arXiv preprint arXiv:1511.07122, 2015.

[38] X. Chen, Y. Yuan, G. Zeng, and J. Wang, “Semi-supervised semantic segmentation with cross pseudo supervision,” 2021.

[39] P. Bachman, O. Alsharif, and D. Precup, “Learning with pseudoensembles,” Advances in neural information processing systems, vol. 27, 2014.

[40] H. Zhang, Z. Zhang, A. Odena, and H. Lee, “Consistency regularization for generative adversarial networks,” in International Conference on Learning Representations, 2019.

[41] L. N. Smith and N. Topin, “Super-convergence: Very fast training of neural networks using large learning rates,” arXiv preprint arXiv:1708.07120, 2017.

[42] G. Ke, Q. Meng, T. Finley, T. Wang, W. Chen, W. Ma, Q. Ye, and T.-Y. Liu, “Lightgbm: A highly efficient gradient boosting decision tree,” Advances in neural information processing systems, vol. 30, 2017.

[43] Y. Ju, G. Sun, Q. Chen, M. Zhang, H. Zhu, and M. U. Rehman, “A model combining convolutional neural network and lightgbm algorithm for ultra-short-term wind power forecasting,” Ieee Access, vol. 7, pp. 28 309–28 318, 2019.

[44] X. Sun, M. Liu, and Z. Sima, “A novel cryptocurrency price trend forecasting model based on lightgbm,” Finance Research Letters, vol. 32, p. 101084, 2020.

[45] G. Liu and J. Guo, “Bidirectional lstm with attention mechanism and convolutional layer for text classification,” Neurocomputing, vol. 337, pp. 325–338, 2019.

[46] W. Li, F. Qi, M. Tang, and Z. Yu, “Bidirectional lstm with self-attention mechanism and multi-channel features for sentiment classification,” Neurocomputing, vol. 387, pp. 63–77, 2020.

[47] A. Vaswani, N. Shazeer, N. Parmar, J. Uszkoreit, L. Jones, A. N. Gomez, Ł. Kaiser, and I. Polosukhin, “Attention is all you need,” Advances in neural information processing systems, vol. 30, 2017.

[48] H. Astola, L. Seitsonen, E. Halme, M. Molinier, and A. Lönnqvist, “Deep neural networks with transfer learning for forest variable estimation using Sentinel-2 imagery in boreal forest,” Remote Sensing, vol. 13, no. 12, 2021.

[49] W. G. Rees, J. Tomaney, O. Tutubalina, V. Zharko, and S. Bartalev, “Estimation of boreal forest growing stock volume in russia from sentinel-2 msi and land cover classification,” Remote Sensing, vol. 13, no. 21, 2021. [Online]. Available: https://www.mdpi.com/2072-4292/13/21/4483

[50] E. Tomppo, O. Antropov, and J. Praks, “Boreal forest snow damage mapping using multi-temporal sentinel-1 data,” Remote Sensing, vol. 11, no. 4, 2019. [Online]. Available: https://www.mdpi.com/2072-4292/11/4/384

[51] W. Huang, W. Min, J. Ding, Y. Liu, Y. Hu, W. Ni, and H. Shen, “Forest height mapping using inventory and multi-source satellite data over hunan province in southern china,” Forest Ecosystems, vol. 9, 2022.

[52] N. Lang, K. Schindler, and J. D. Wegner, “Country-wide high-resolution vegetation height mapping with sentinel-2,” Remote Sensing of Environment, vol. 233, p. 111347, 2019.

[53] H. Astola, T. Häme, L. Sirro, M. Molinier, and J. Kilpi, “Comparison of sentinel-2 and landsat 8 imagery for forest variable prediction in boreal region,” Remote Sensing of Environment, vol. 223, pp. 257–273, 2019. [Online]. Available: https://www.sciencedirect.com/science/article/pii/S0034425719300252

[54] J. Praks, O. Antropov, and M. T. Hallikainen, “Lidar-aided sar interferometry studies in boreal forest: Scattering phase center and extinction coefficient at x-and l-band,” IEEE Transactions on Geoscience and Remote Sensing, vol. 50, no. 10 PART1, p. 3831–3843, 2012.

[55] A. Olesk, J. Praks, O. Antropov, K. Zalite, T. Arumäe, and K. Voormansik, “Interferometric sar coherence models for characterization of hemiboreal forests using tandem-x data,” Remote Sensing, vol. 8, no. 9, 2016.

[56] F. Kugler, D. Schulze, I. Hajnsek, H. Pretzsch, and K. P. Papathanassiou, “Tandem-x pol-insar performance for forest height estimation,” IEEE Transactions on Geoscience and Remote Sensing, vol. 52, no. 10, p. 6404–6422, 2014.

